# Integration of GWAS and TWAS to elucidate the genetic architecture of natural variation for leaf cuticular conductance in maize

**DOI:** 10.1101/2021.10.26.465975

**Authors:** Meng Lin, Pengfei Qiao, Susanne Matschi, Miguel Vasquez, Guillaume P. Ramstein, Richard Bourgault, Marc Mohammadi, Michael J. Scanlon, Isabel Molina, Laurie G. Smith, Michael A. Gore

## Abstract

The cuticle, a hydrophobic layer of cutin and waxes synthesized by plant epidermal cells, is the major barrier to water loss when stomata are closed. Dissecting the genetic architecture of natural variation for maize leaf cuticular conductance (*g*_c_) is important for identifying genes relevant to improving crop productivity in drought-prone environments. To this end, we performed an integrated genome- and transcriptome-wide association study (GWAS/TWAS) to identify candidate genes putatively regulating variation in leaf *g*_c_. Of the 22 plausible candidate genes identified, five were predicted to be involved in cuticle precursor biosynthesis and export, two in cell wall modification, nine in intracellular membrane trafficking, and seven in the regulation of cuticle development. A gene encoding an INCREASED SALT TOLERANCE1-LIKE1 (ISTL1) protein putatively involved in intracellular protein and membrane trafficking was identified in GWAS and TWAS as the strongest candidate causal gene. A set of maize nested near-isogenic lines that harbor the *ISTL1* genomic region from eight donor parents were evaluated for *g*_c_, confirming the association between *g*_c_ and *ISTL1* in a haplotype-based association analysis. The findings of this study provide novel insights into the role of regulatory variants in the development of the maize leaf cuticle, and will ultimately assist breeders to develop drought-tolerant maize for target environments.

**Sentence summary:** We performed an integrated GWAS/TWAS and identified 22 candidate genes putatively regulating variation in maize leaf *g*_c_. The association between *g*_c_ and the strongest candidate causal gene, *ISTL1*, was validated with maize nested near-isogenic lines.

## Introduction

The cuticle is a hydrophobic layer covering the epidermal surfaces of the shoot, which protects plants from dehydration, UV radiation, and pathogen attack (Shepherd and Wynne Griffiths, 2006; Xue et al., 2017). Cutin and waxes are the two major lipid components comprising the plant cuticle. Cutin is an insoluble matrix formed by extensive ester cross-linking of fatty acid derivatives and glycerol (Pollard et al., 2008; Fich et al., 2016). Soluble waxes, including alkanes, aldehydes, alcohols, ketones, and wax esters, are embedded within and on top of the cutin matrix (Yeats and Rose, 2013).

Cuticle composition and structure influence its water barrier function (Kerstiens, 2006). However, cuticle impermeability to water is not simply determined by wax load or cuticle thickness; instead, chemical composition and the organization of cuticle components appear to contribute to reducing non-stomatal water loss (Riederer and Schreiber, 2001). Thus, cuticle composition and structure are potentially relevant to plant drought tolerance. Mutants and transgenic plants with reduced wax load and compromised cuticle structure often show increased cuticular permeability and decreased drought tolerance (Zhou et al., 2013; Zhu and Xiong, 2013; Li et al., 2019), while overexpression of cuticle lipid biosynthesis enzymes or their transcriptional regulators can result in increased cuticular lipid abundance and drought tolerance (Aharoni et al., 2004; Zhang et al., 2005; Bourdenx et al., 2011; Wang et al., 2012). In wheat and barley, glaucousness, a visual manifestation of epicuticular wax crystals, was selected for during domestication (Bi et al., 2016; Hen-Avivi et al., 2016) and shows positive correlations with drought tolerance (Febrero et al., 1998; Guo et al., 2016; Busta et al., 2021). Collectively, these findings suggest a range of possibilities toward increased drought tolerance via cuticle modification in cereal crops.

Biological pathways involved in cuticle development affect its permeability to water in different ways. The formation of the plant cuticle requires not only the synthesis of cutin monomers and cuticular waxes but also the transport and polymerization of cutin monomers after transport (Yeats and Rose, 2013). Major pathways for the biosynthesis of cutin monomers and cuticular waxes have been elucidated in model plant systems (Yeats and Rose, 2013; Lee and Suh, 2015; Fich et al., 2016). A Golgi-mediated vesicle trafficking system delivers cuticle lipids from the intracellular membranes, where they are synthesized, to the plasma membrane (McFarlane et al., 2014) for export by ATP BINDING CASSETTE TRANSPORTER G (ABCG) family proteins (Pighin et al., 2004; Bird et al., 2007; Panikashvili et al., 2010; Bessire et al., 2011; Chen et al., 2011) and extracellular lipid transport proteins (DeBono et al., 2009; Kim et al., 2012). In addition, GLY-ASP-SER-LEU ESTERASE/LIPASE (GDSL lipase) is required for cutin polymerization in the developing cuticle (Girard et al., 2012; Yeats and Rose, 2013), and the degree of pectin esterification is critical to cuticle structure by facilitating the diffusion of cutin precursors (Bakan and Marion, 2017; Philippe et al., 2020). Transcription factors have also been identified to regulate various steps in cuticle development (Borisjuk et al., 2014; Elango et al., 2020), but how these transcription factors, biosynthesis and transport processes work together is not well understood.

Genome-wide association studies (GWAS) are widely used to identify genomic variants that are associated with a trait of interest (Tibbs Cortes et al., 2021). A genome-wide association study conducted by Lin et al. (2020) revealed that adult maize leaf cuticular conductance (*g*_c_; defined as the rate of water loss when stomata are closed, an indirect measure of cuticular permeability to water) is likely controlled by large numbers of small effect alleles in maize, which agrees with the complex cuticle biosynthesis network and transport mechanisms (Yeats and Rose 2013). Given that the GWAS of Lin et al. (2020) was limited by low mapping resolution and statistical power, the use of gene expression data with its offering of gene-level resolution and capturing of regulatory variation provides an opportunity to address such weaknesses. Therefore, transcriptome-wide association studies (TWAS), which identify significant associations between trait variation and transcript abundance across all expressed genes in a tissue, can provide complementary information for candidate gene identification in an independent test. Furthermore, integrating GWAS and TWAS results with an ensemble approach based on the Fisher’s combined test (FCT) improves statistical power in candidate gene identification (Kremling et al., 2019), especially for leaf physiological traits with a complex genetic architecture (Ferguson et al., 2021; Pignon et al., 2021).

In this study, an integrated investigation was performed to dissect natural variation for leaf *g*_c_ in a maize inbred association panel scored with ∼9 million SNP markers and transcript abundance from cuticle maturation zones (as identified by Bourgault et al. 2020) of adult leaves. Combining GWAS and TWAS results with a FCT boosted statistical power to identify genes for this complex trait. Evaluation of maize nested near-isogenic lines (nNILs) (Morales et al., 2020) confirmed the association between *g*_c_ and one of the identified plausible causal genes, which encodes a predicted INCREASED SALT TOLERANCE1-LIKE1 (ISTL1) protein.

## Results

### Relationship of *g*_c_ to cuticle composition and structure

Permeability of the leaf cuticle to water was measured as *g*_c_ on 310 genetically representative maize inbred lines from the Wisconsin Diversity (WiDiv) panel that were evaluated in two environments in San Diego, CA. To investigate how this phenotype of moderately high heritability (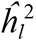= 0.73) related to cuticle characteristics, we quantitatively analyzed selected features of adult leaf cuticle composition and structure in a subset of 51 WiDiv lines with highest and lowest *g*_c_ values. Our structural analysis focused on features of guard cell cuticles that could be analyzed quantitatively in a large number of samples via confocal microscopy, and seemed likely to affect the *g*_c_ trait. As illustrated in Supplemental Figure 1, we observed cuticular flaps projecting from the inner surfaces of guard cell pores, as described in many plant species, often called “outer cuticular ledges” (e.g. Hunt et al., 2017), which have been proposed to reduce water loss by enhancing the seal of closed stomata (Wilmer and Fricker, 1996; Zhao and Sack, 1999; Kosma and Jenks, 2007). We also observed cuticular bumps lining stomatal pores, which appear to permit a “zippering” of closed stomata (Supplemental Figure 1). However, variation in these traits was only weakly associated with *g*_c_ (⎟*r*⎟ < 0.15 for all measures; Supplemental Table S1). Numerous studies have demonstrated relationships between cuticular wax composition and cuticle permeability (Zhou et al., 2013; Zhu and Xiong, 2013; Li et al., 2019), and wax composition was also found to vary in these extreme *g*_c_ lines. Therefore, we also investigated how variation in *g*_c_ may be related to wax composition. Significant associations were observed between *g*_c_ and a subset of fatty acids, wax esters, aldehydes, and alicyclics (*r* = −0.299 to −0.488; Supplemental Table S1).

To further query the relationship of *g*_c_ to cuticular wax composition and structural features, random forest analysis was conducted with five-fold cross-validation to determine the degree to which *g*_c_ can be predicted by the analyzed features, and which features are the strongest drivers of predictive ability. As shown in Figure 1 and Supplemental Table S1, we found that the abundance of several long chain wax esters and aldehydes (especially C54 wax esters and C30 aldehydes) to be important predictors of *g*_c_. The overall predictive ability for *g*_c_ from all measured characteristics combined was 0.35, with most of the predictive power coming from measurements of C54 wax esters and C30 aldehydes. The relationship of *g*_c_ to C54 wax ester abundance is interesting in relation to the findings of Bourgault et al. (2020), showing that acquisition of mature water barrier function during cuticle development in adult maize leaves coincided with a shift from alkanes to esters with carbon chain lengths > 49 as the most abundant wax class. Thus, our analysis of wax composition in extreme lines provides further evidence of a role for wax esters in protecting adult maize leaves against water loss, consistent with recent findings in Arabidopsis (Patwari et al., 2019). However, the majority of *g*_c_ is not explained by the cuticle features we analyzed, underscoring the complexity of this trait that merits further genetic examination through the combination of GWAS and TWAS.

**Figure 1.**
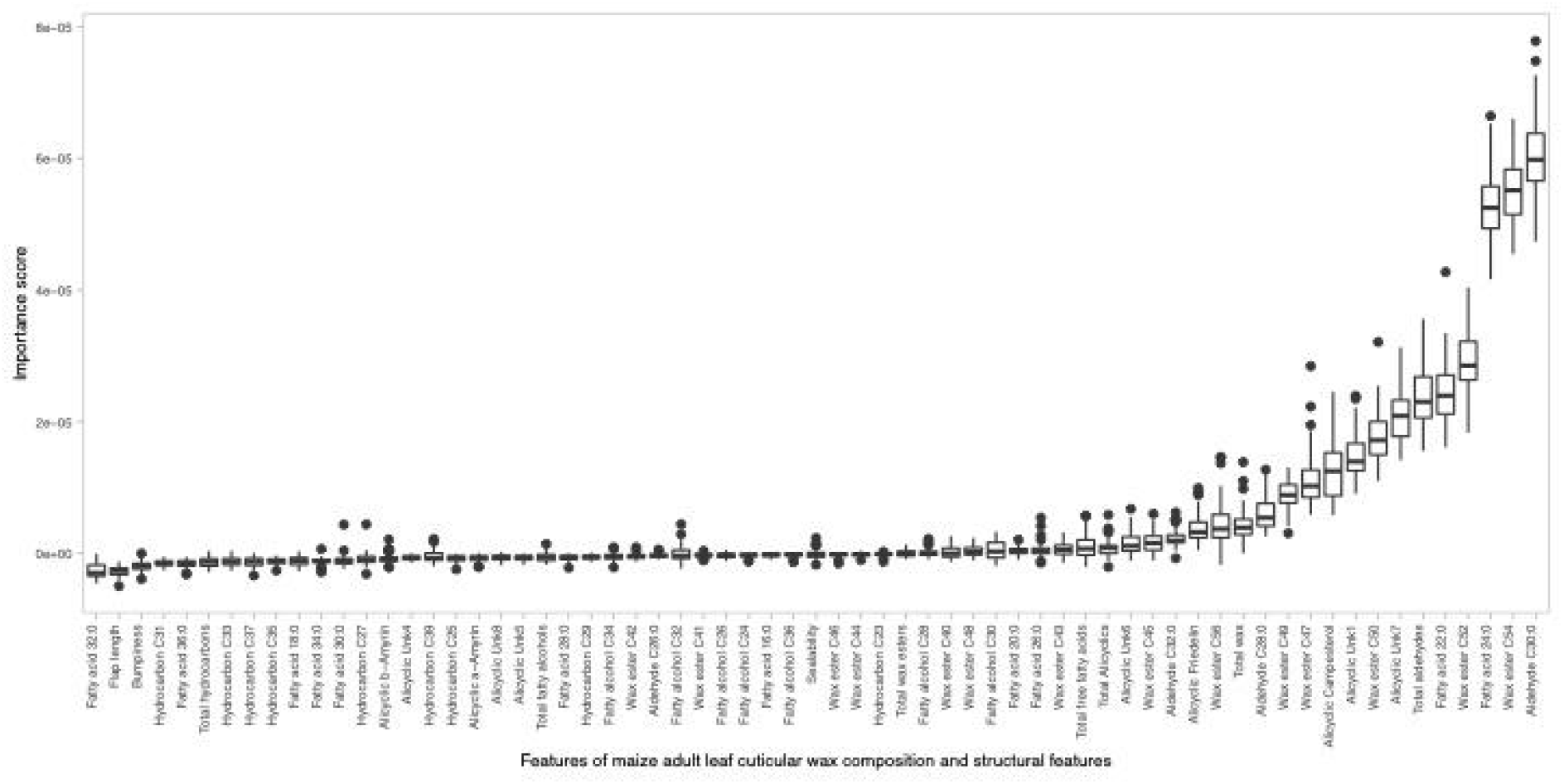
Box-plot of importance scores for cuticular wax composition and structural features in random forest regression to predict maize leaf cuticular conductance (*g*_c_) using five-fold cross validation for 50 times.

### Quantitative genetic analysis for identification of candidate genes for *g*_c_

To improve statistical power in the identification of candidate genes for *g*_c_ beyond that achieved in GWAS alone by Lin et al. (2020), we performed the Fisher’s combined test (FCT) to integrate GWAS and TWAS results (see Methods for more detail). A total of 22 plausible candidate genes were selected based on predicted functions that could be connected to cuticles from a total of 319 candidates (top 1%) from FCT (Supplemental Table S2), 273 (top 0.01%) from GWAS (Supplemental Tables S3 and S4), and 200 (top 1%) from TWAS (Supplemental Table S5). Of these 22 genes, 14 were identified by FCT, five by TWAS and three by GWAS; nine were identified by two or more of these strategies (Table 1; Supplemental Figure S2). Supplemental Table S6 lists the closest Arabidopsis and rice relatives of these 22 genes.

**Table 1.** Plausible candidate genes identified in a genome-wide association study (GWAS), a transcriptome-wide association study (TWAS) and the Fisher’s combined test (FCT) for leaf cuticular conductance (*g*_c_) in the Wisconsin diversity panel.

| Gene ID | Annotation | Chromosome | Gene start | Gene end | Functional category<sup>b</sup> | Method |
| --- | --- | --- | --- | --- | --- | --- |
| Zm00001d029141 | MIEL1-like | 1 | 60,322,950 | 60,326,019 | RCD | GWAS, FCT |
| Zm00001d032788 | GDSL esterase/lipase | 1 | 238,190,770 | 238,192,464 | CBE | TWAS, FCT |
| Zm00001d002843 | MYB54 | 2 | 24,293,846 | 24,295,521 | RCD | FCT |
| Zm00001d005087 | GLK35 | 2 | 157,736,007 | 157,737,776 | RCD | TWAS, FCT |
| Zm00001d039411 | α1-COP | 3 | 3,708,911 | 3,716,373 | IMT | TWAS, FCT |
| Zm00001d040788 | SEC15B | 3 | 65,988,895 | 65,991,327 | IMT | GWAS |
| Zm00001d042612 | CER7 | 3 | 173,571,381 | 173,573,177 | RCD | FCT |
| Zm00001d043509 | PME40 | 3 | 202,333,699 | 202,335,784 | CWB | TWAS |
| Zm00001d044162 | WRKY64 | 3 | 220,824,499 | 220,827,409 | RCD | TWAS |
| Zm00001d048953 | KCS3 | 4 | 10,603,315 | 10,607,553 | CBE | TWAS, FCT |
| Zm00001d049479 | ISTL1 | 4 | 31,729,260 | 31,732,664 | IMT | GWAS, TWAS, FCT |
| Zm00001d050185 | MYB108 | 4 | 71,478,007 | 71,479,474 | RCD | TWAS |
| Zm00001d051599 | Rab-GAP/TBC | 4 | 164,071,426 | 164,090,678 | IMT | TWAS, FCT |
| Zm00001d012964 | SEC14 | 5 | 2,431,015 | 2,437,696 | IMT | FCT |
| Zm00001d015477 | CER3-like | 5 | 92,669,605 | 92,675,745 | CBE | TWAS, FCT |
| Zm00001d036765 | CER9 | 6 | 100,099,887 | 100,107,565 | RCD | TWAS |
| Zm00001d038404/<br>Zm00001d038405<sup>a</sup> | Ypt/Rab-GAP | 6 | 156,394,901/<br>156,409,880 | 156,403,240/<br>156,412,505 | IMT | GWAS |
| Zm00001d022364 | SNARE | 7 | 176,149,978 | 176,160,500 | IMT | TWAS, FCT |
| Zm00001d010426 | CER5/ABCG12-like | 8 | 114,527,255 | 114,533,597 | CBE | TWAS |
| Zm00001d012175 | PAE5 | 8 | 169,547,499 | 169,551,527 | CWB | FCT |
| Zm00001d025180 | Ypt/Rab-GAP | 10 | 107,999,211 | 108,003,428 | IMT | FCT |

<sup>a</sup>In B73 RefGen\_v4, this gene was erroneously split into the two gene models as indicated, but in B73 RefGen\_v5 they have been correctly merged into a single gene model with identifier Zm00001eb289920.
<sup>b</sup>RCD, regulator of cuticle development; CBE, cuticle biosynthesis and export; IMT, intracellular membrane trafficking; CWB, cell wall biosynthesis.

Of the 14 genes identified by FCT, one (*Zm00001d049479*) encoding the closest maize relative of INCREASED SALT TOLERANCE1-LIKE1 (ISTL1) in Arabidopsis, which functions in multivesicular body formation within endosomes (Buono et al,. 2016) was also identified by both TWAS and GWAS, and was the only candidate gene identified by all three methods (Table 1; Figure 2). Additionally, five of the 14 genes identified by FCT encode proteins with predicted functions in intracellular membrane trafficking, three of which were also identified by TWAS (Table 1). Two of these five, *Zm00001d039411* and *Zm00001d012964*, encode proteins with sequence identity to α1 COAT PROTEIN (α1-COP) and SEC14 in Arabidopsis, respectively, which function in vesicle formation at the Golgi surface (Jouannic et al., 1998; Lee and Goldberg, 2010). *Zm00001d051599* and *Zm00001d025180* encode proteins with Rab GTPase-activating protein (Rab-GAP) domains, predicted to function in regulation of vesicle transport (Stenmark, 2009), and *Zm00001d022364* encodes a soluble NSF attachment protein receptor (SNARE) homolog with an expected function in vesicle fusion with target membranes (Chen and Scheller, 2001).

**Figure 2.**
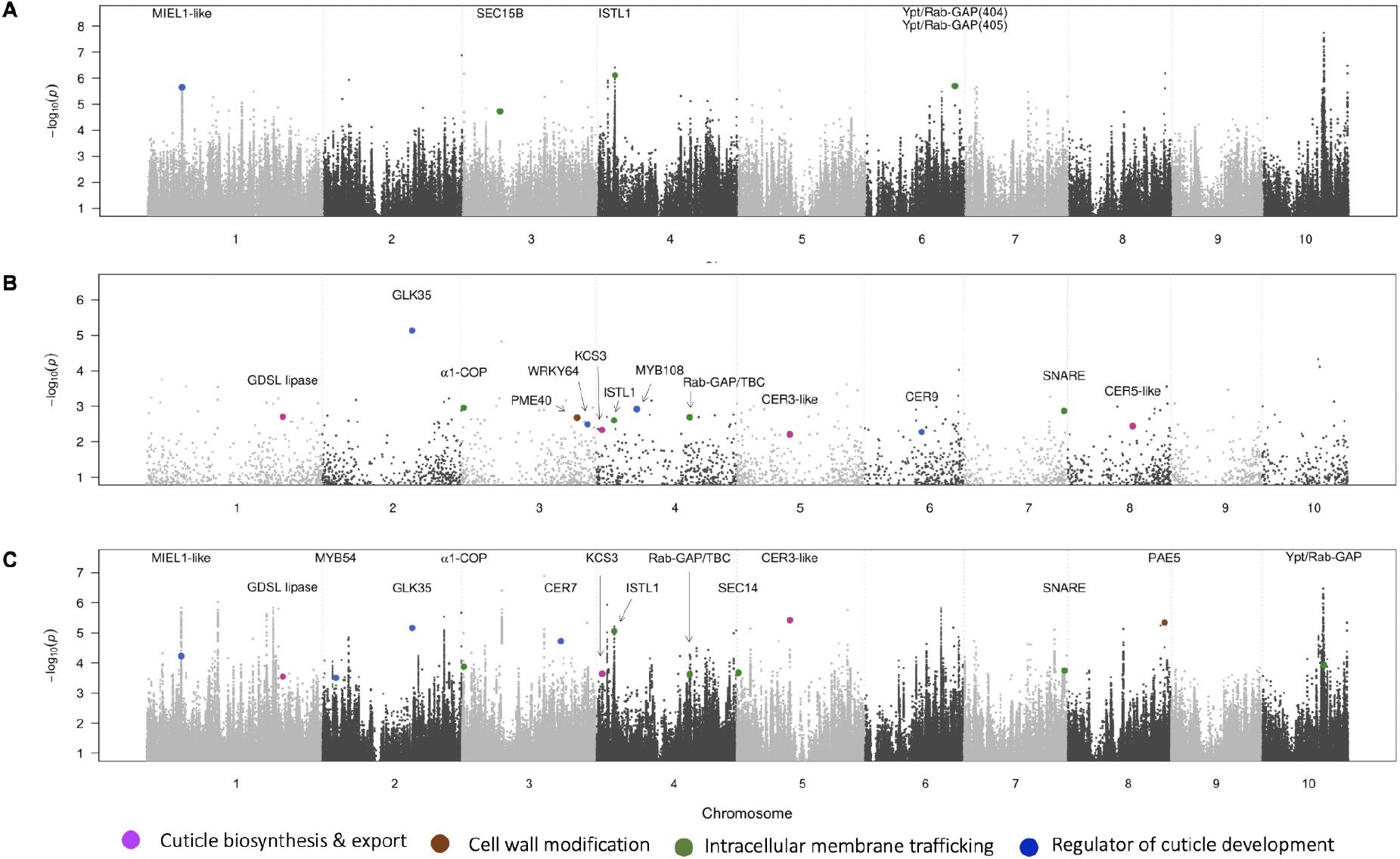
Manhattan plot of results from a genome-wide association study (A; GWAS), a transcriptome-wide association study (B; TWAS), and Fisher’s combined test (C; FCT) of adult maize leaf cuticular conductance (*g*_c_) across two environments in San Diego, CA. (A) The -log_10_ *P*-value of each SNP tested in a mixed linear model analysis of *g*_c_ is plotted as a point against its physical position (B73 RefGen_v4) for the 10 chromosomes of maize. The peak SNPs closest to plausible candidate genes are colored based on their functional categories. (B) The -log_10_ *P*-value of each gene transcript tested in a mixed linear model analysis of *g*_c_ is plotted as a point against its physical position (B73 RefGen_v4). Plausible candidate genes are colored based on their functional categories. (C) The -log_10_ *P*-value of each of the top 10% SNPs in GWAS paired with the nearest gene in FCT plotted as a point against its physical position (B73 RefGen_v4). The most significant SNP-gene pairs related to plausible candidate genes identified in the FCT are colored based on their functional categories.

The remaining eight plausible candidate genes identified by FCT, four of which were also identified by TWAS and one also by GWAS, encode proteins with a variety of functions. Two (*Zm00001d048953* and *Zm00001d015477*, *3-KETOACYL-COA SYNTHASE 3* [*KCS3*] and *GLOSSY1-like/ECERIFERUM3-like* [*GL1-like/CER3-like*], respectively), are homologs of Arabidopsis genes with functions in cuticular wax biosynthesis (Lee et al., 2009; Bernard et al., 2012). A third (*Zm00001d032788*) encodes a predicted GDSL esterase/lipase gene closely related to Arabidopsis and tomato cutin synthases (Girard et al., 2012; Yeats et al., 2012). Two FCT-identified genes (*Zm00001d005087* and *Zm00001d002843,* annotated as GLK35 and MYB54, respectively), encode transcription factors with potential functions in regulation of cuticle biosynthesis (Kant et al., 2008; Dubos et al., 2010; Oshima and Mitsuda, 2013). *Zm00001d029141* is a homolog of Arabidopsis MIEL1 (MYB30-INTERACTING E3 LIGASE 1), which regulates the stability of two MYB family regulators of wax biosynthesis (Marino et al., 2013; Gil et al., 2017). *Zm00001d042612* encodes a homolog of Arabidopsis CER7, a core subunit of the RNA degrading exosome associated with alkane biosynthesis (Hooker et al., 2007). The final FCT-identified gene listed in Table 1, *Zm00001d012175* (PAE5), encodes a predicted pectin-modifying enzyme.

A total of five additional plausible candidate genes were uniquely detected by TWAS (Figure 2; Table 1). *Zm00001d043509* encodes a putative pectin methylesterase (PME40). *Zm00001d010426* encodes a homolog of Arabidopsis CER5/ABCG12, which functions in cuticular wax secretion (Pighin et al., 2004). *Zm00001d036765* encodes a homolog of Arabidopsis CER9, which regulates the abundance of both cutin and cuticular waxes (Lü et al., 2012). The two remaining TWAS-identified genes (*Zm00001d050185* and *Zm00001d044162*) encode homologs of Arabidopsis WRKY and MYB family transcription factors with potential roles in cuticle development, which are differentially expressed across maize leaf cuticular developmental stages (Qiao et al., 2020).

GWAS uniquely identified three additional plausible candidate genes predicted to be involved in the intracellular membrane trafficking system (Figure 2; Table 1). Two adjacent genes (*Zm00001d038404* and *Zm00001d038405*) putatively encode Ypt/Rab-GAPs, regulators of vesicle fusion (Stenmark, 2009). *Zm00001d040788* encodes a homolog of Arabidopsis SEC15B, an exocyst subunit that activates vesicle fusion with the plasma membrane during polarized secretion (Guo et al., 1999; Mayers et al., 2017).

### Haplotype-based association analysis for *g*_c_ in the WiDiv panel

The *ISTL1* gene emerged from our analysis as a strong candidate regulator of *g*_c_, since it was identified in our prior GWAS (Lin et al., 2020) and the only gene detected by all three analyses (GWAS, TWAS and FCT) in this study. To further investigate the association between *g*_c_ and genetic variation within the WiDiv population in the vicinity of *ISTL1*, we analyzed the association between *g*_c_ and haplotypes at 91 haploblocks (blocks of haplotypes) within 200 kb of *ISTL1.* Of these, Block 65 was the most significantly associated with *g*_c_ (*P*-value 4.07 × 10^-7^) in the WiDiv panel (Figure 3; Supplemental Table S7), having three haplotypes defined by three SNPs including the peak GWAS SNP (4-30231047) only 1,219 bp away from the 3′ end of *ISTL1*. The second most strongly associated (*P*-value 4.07 × 10^-7^) haploblock, Block 52 (Figure 3; Supplemental Table S7), consisted of 10 haplotypes, with the 16 defining SNPs spanning a 8.6 kb region more than 20 kb upstream of *ISTL1*. These findings suggest that the underlying causal variants are located both upstream and downstream of the *ISTL1* gene.

**Figure 3.**
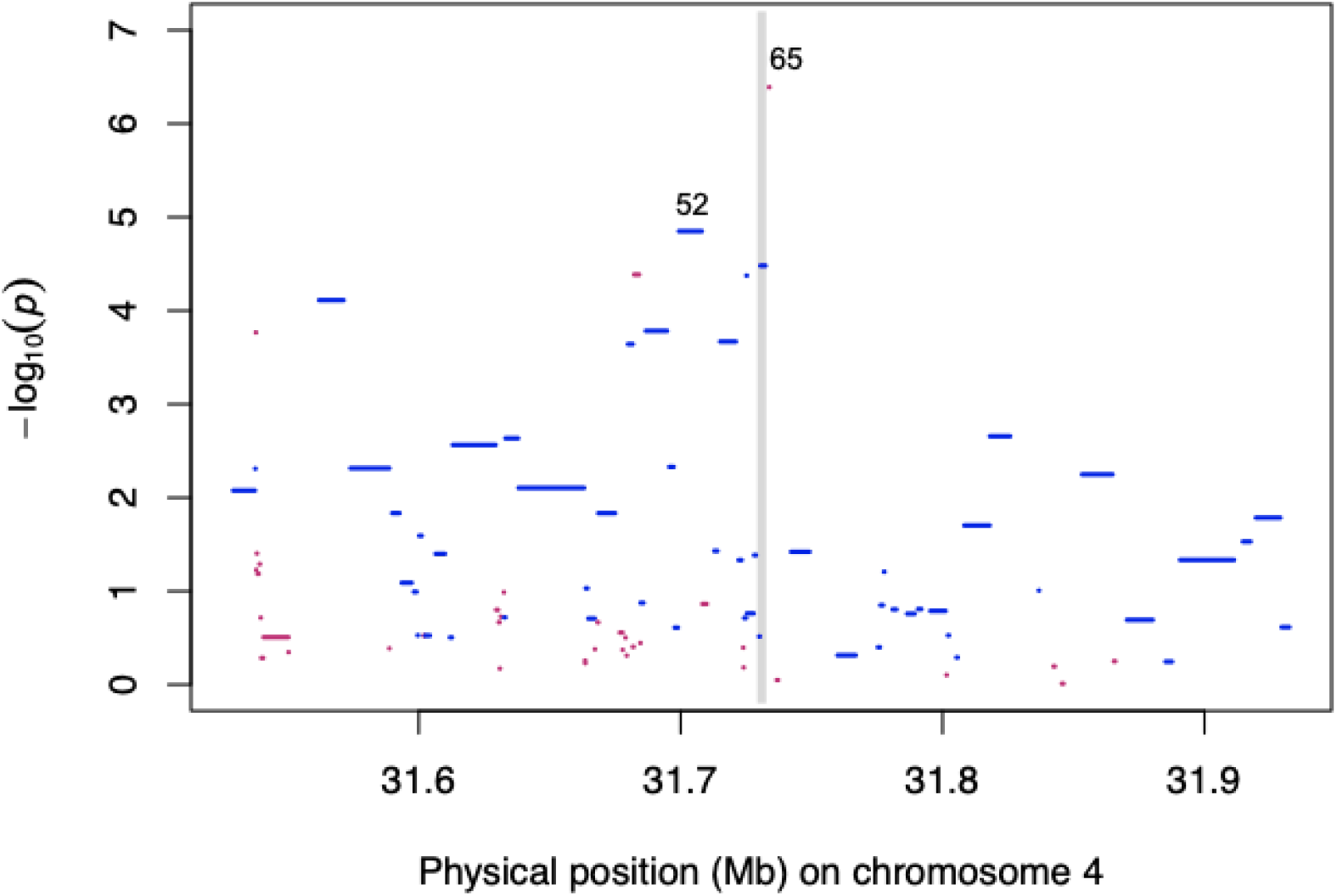
Manhattan plot of results from a haplotype-based association analysis in the vicinity of *ISTL1* for adult maize leaf cuticular conductance (*g*_c_) combined from two environments in San Diego, CA, in the Wisconsin diversity panel. The -log_10_ *P*-value of each haploblock tested in a mixed linear model analysis of *g*_c_ is plotted as a short horizontal line against its physical position (B73 RefGen_v4). The length of a short horizontal line corresponds to the size of that haplotype block. The grey vertical bar represents the physical position of *ISTL1*. The top two haploblocks (Blocks 65 and 52) that are most associated with *g*_c_ are indicated. Polymorphic and monomorphic haploblocks among the evaluated nested near-isogenic lines are colored in blue and red, respectively.

### Validation of *ISTL1* haplotype effects with nested NILs

A set of 12 nested near-isogenic lines (nNILs) (Morales et al., 2020) containing the *ISTL1* genomic region from diverse inbreds that are mostly not represented in the WiDiv panel, introgressed into B73 (Supplemental Table S8), were used to validate haplotype effects observed at *ISTL1* in the WiDiv panel. All 12 evaluated nNILs had lower *g*_c_ relative to their corresponding donor parent (Figure 4; Supplemental Table S9), which is to be expected given that the recurrent parent, B73, had a comparable or lower *g*_c_ than the donor parents. Compared to B73, two NILs with Tzi8 introgressions (Tzi8-B73_NIL_1160 and Tzi8-B73_NIL_1308) and one NIL with an Oh43 introgression (Oh43-B73_NIL_1005) had significantly lower *g*_c_, while four of the donor parents (Ki11, NC350, Oh43 and Tx303) showed significantly higher *g*_c_ (Figure 4; Supplemental Table S9).

**Figure 4.**
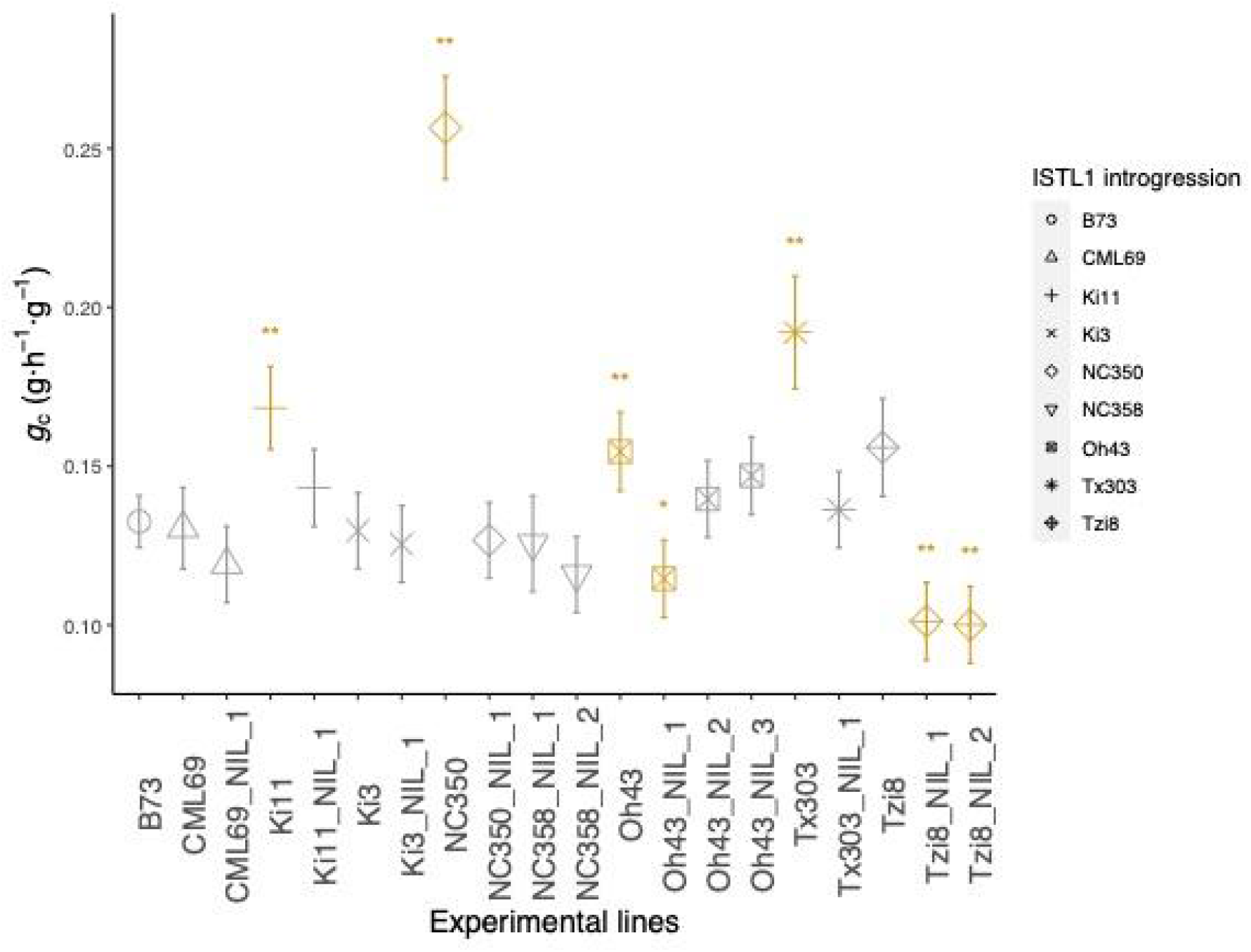
Scatter plot of best linear unbiased estimator values for adult maize leaf cuticular conductance (*g*_c_) for B73, nested near-isogenic lines (nNILs) containing *ISTL1* introgressions and their donor parent lines across two environments in San Diego, CA, in 2020. Lines with *g*_c_ significantly different from B73 are highlighted in orange. ‘*’ represents *P*-value < 0.1 and ‘**’ represents *P*-value < 0.05 after adjustment by the Dunnett’s method.

Of the two most strongly associated haploblocks in the WiDiv panel, only Block 52 (Supplemental Figure S3A), the second ranked haplotype-based association, was polymorphic among the nNILs (Figure 3). This haploblock, consisting of three segregating haplotypes (Supplemental Figure S3B), had a significant association (*P*-value < 0.020) with *g*_c_ among nNILs. The haplotype effect of the Tzi8 NILs, which possess a unique haplotype relative to B73 and the other donor parents, was significantly (*P*-value < 0.05) different from the other two haplotypes (Supplemental Figures S3B). The Tzi8 haplotype had an effect of 0.1007 g·h^-1^·g^-1^, whereas the other two haplotypes averaged ∼0.13·h^-1^·g^-1^. The smaller effect of the Tzi8 haplotype could explain why both Tzi8 nNILs had the lowest observed *g*_c_ values (Figure 4). Haplotype effects of Block 52 strongly correlated (*r* = 0.924) with those estimated in the WiDiv panel (Supplemental Figure S3C), indicating that the haplotype effects at *ISTL1* for *g*_c_ could be validated in an independent panel having a different population structure.

## Discussion

The plant cuticle, as a crucial barrier to water loss (Shepherd and Wynne Griffiths, 2006; Xue et al., 2017), is potentially related to plant drought tolerance (Aharoni et al., 2004; Wang et al., 2012; Zhou et al., 2013; Li et al., 2019). Although major pathways for wax and cutin monomers biosynthesis have been elucidated in model species (Yeats and Rose, 2013; Bakan and Marion, 2017), genetic controls of water loss through leaf cuticles, i.e. *g*_c_, remain largely unknown. In a prior study (Lin et al., 2020), we used 235,004 GBS SNP markers for a GWAS utilizing 451 lines of the WiDiv panel to identify five genomic regions impacting *g*_c_. In the present study, we extended this earlier work by imputing genotypes for ∼9 million SNPs and evaluating the following two features for a subset of 310 WiDiv lines in two growth environments: *g*_c_, and transcript abundance for 20,013 genes in the cuticle maturation zones of developing adult leaf blades. These data were analyzed by GWAS, TWAS, and FCT, greatly increasing the power to identify genes with likely roles in determination of *g*_c_. A total of 22 high-scoring genes (discussed below) from one or more of these tests were identified as plausible candidates based on predicted functions potentially impacting *g*_c_ via the regulation of cell wall biosynthesis (2 genes), cuticle biosynthesis and export (4 genes), intracellular membrane trafficking (9 genes) and cuticle development (7 genes) (Figure 2; Table 1). Of the 23 GWAS peaks detected in this study, only the GWAS signal associated with *ISTL1* was also found in our previous study that identified five genomic loci associated with *g*_c_ ^(^Lin et al., 2020^)^. The main reason for such limited overlap is most likely the extreme difference in the growth environment where associations were found in the prior study (Maricopa, AZ which is very hot and dry) compared to that used for the present study (San Diego, CA, which is relatively cool and humid). Subtle changes in population structure when subsetting the WiDiv population, and different criteria for calling GWAS peaks in this study, may also have contributed to differences in GWAS signals.

Two of the genes identified as plausible candidates for *g*_c_ determination have predicted functions in cell wall modification, in particular pectin esterification (PAE5 and PME40). Cutin-embedded polysaccharides present a high degree of esterification, including methylation and acetylation (Philippe et al., 2020). Pectin esterification could be critical to cuticle structure by favoring interactions between cutin and cutin-embedded polysaccharides as well as facilitating the diffusion of cutin precursors (Bakan and Marion, 2017; Philippe et al., 2020) potentially modulating cuticle function as a barrier to water loss.

Our study also pointed to genes encoding predicted cuticle lipid biosynthetic enzymes. FCT identified a maize homolog of *CUS1/LTL1*, encoding a GDSL acyltransferase that polymerizes cutin precursors extracellularly in Arabidopsis and tomato (Girard et al., 2012; Yeats et al., 2012). FCT and TWAS both identified maize KCS3, a putative component of the fatty acid elongase (FAE) complex that extends acyl chains produced by the plastid to very-long-chain fatty acids. Its closest Arabidopsis relative, KCS2, functions in the two-carbon elongation of acyl chains to C22 very long chain fatty acyl precursors required for cuticular wax and root suberin biosynthesis (Franke et al., 2009; Lee et al., 2009). Notably, several *KCS* homologs are upregulated during cuticle maturation in the developing maize adult leaf epidermis (Qiao et al., 2020). TWAS and FCT also identified a maize homolog of *AtCER3* and *ZmGL1*, genes involved in alkane biosynthesis (Sturaro et al., 2005; Bernard et al., 2012). However, this *GL1/CER3*-like maize gene has not been previously characterized by forward or reverse genetic approaches. Finally, TWAS analysis identified a maize homolog of Arabidopsis *ABCG12/CER5*, encoding a half-transporter of the ABCG family that forms heterodimers with ABCG11 to transport cuticle lipids across the plasma membrane (Pighin et al., 2004; McFarlane et al., 2010). Unlike the other cuticle biosynthesis candidates highlighted in this study, this maize *CER5* homolog is epidermally upregulated during cuticle maturation (Qiao et al., 2019). Identification of several candidate genes with predicted functions in the biosynthesis and transport of cuticle lipids is consistent with our finding that *g*_c_ can be partially predicted by cuticular wax composition.

The biosynthesis of plant cuticles is regulated in a complex network (Yeats and Rose, 2013). In this study, we identified maize homologs of three known regulators of cuticle development (Figure 2; Table 1). This includes a maize homolog of Arabidopsis CER7, a core subunit of exosomal 3′-to-5′ exoribonuclease. This enzyme reduces the degradation of CER3/WAX2/YRE transcripts (Hooker et al., 2007). CER3 and CER1 proteins interact to jointly synthesize VLC-alkane (Bernard et al., 2012), an important class of cuticle waxes that is related to drought resistance (Kosma et al., 2009; Wang et al., 2015; Wang et al., 2020). Interestingly, a different maize CER7 homolog was associated with *g*_c_ in our prior GWAS (Lin et al., 2020), underscoring the potential importance of CER7 proteins in the regulation of *g*_c_. TWAS identified maize CER9, homologous to an Arabidopsis E3 ubiquitin ligase whose deficiency enhances plant drought tolerance via alteration of cuticle composition and reduction of cuticular transpiration (Lü et al., 2012). Finally, both GWAS and FCT identified a maize homolog of Arabidopsis MIEL1, an E3 ligase that regulates ABA sensitivity by controlling the protein stability of MBY96 and MYB30, R2R3-MYB subfamily regulators of extracellular lipid biosynthesis (Raffaele et al., 2008; Seo et al., 2011), which are important for balanced cuticular wax biosynthesis (Marino et al., 2013; Lee and Seo, 2016; Gil et al., 2017).

Four putative transcriptional regulators of cuticle biosynthesis were also identified as plausible *g*_c_ candidates: one WRKY, one homeodomain, and two MYB transcription factors (Figure 2; Table 1). *WRKY64* has not been directly associated with cuticle development, but a closely related gene in Arabidopsis (*AtWRKY28; mybAT4G18170*) (Supplemental Table S6) (Jiang et al., 2014) and its maize homolog, *ZmWRKY28* (*Zm00001d011413*), are implicated in cuticle development (Qiao et al., 2020). MYB108 is homologous to ATMYB36, which directly and positively regulates formation of the Casparian strip (characterized by deposition of suberin, which is chemically similar to cutin) (Kamiya et al., 2015). MYB108 is also differentially expressed across the time course of maize leaf cuticular maturation (Qiao et al., 2020). The closest Arabidopsis relative (ATMYB17) of FCT-identified MYB54 (Supplemental Table S6) belongs to the same clade as two well-characterized MYBs of cuticle biosynthesis, MYB106 and MYB16 (Dubos et al., 2010; Oshima and Mitsuda, 2013). GLK35, identified via TWAS and FCT, is a predicted homeodomain-containing transcription factor that has not been previously studied, but its Arabidopsis homolog (AT2G40260) showed > 8-fold change in transcript abundance in response to heat stress (Kant et al., 2008). Therefore, GLK35 could be a regulator for *g*_c_ induced by high temperatures. Together, these candidate genes highlight the potential for developmental and environmental regulation of cuticle formation through transcriptional, post-transcriptional, and post-translational mechanisms.

Several candidate genes identified in our study have predicted functions in intracellular membrane trafficking, the process by which proteins and lipids are moved from one cellular location to another. These candidates are noteworthy because of prior evidence of a role for membrane trafficking in cuticle formation (McFarlane et al., 2014). Two of our candidate genes encode proteins predicted to function in vesicle formation.

One of these is a putative COPI coat component (α1-COP). COPI has not been widely studied in plants, but COPI deficiency in yeast leads to protein accumulation in the ER and global secretion deficiency (Lodish et al., 2016). The other is a putative member of SEC14 family phosphatidylinositol transfer protein, and is different from the two SEC14 homologs we identified previously as candidate regulators of *g*_c_ via GWAS (Lin et al., 2020). Four of our candidate genes have predicted functions in vesicle targeting or fusion, including one putative SNARE, a putative exocyst subunit (SEC15B), and three Rab-GAP domain family proteins with expected functions in regulation of Rab GTPases. By different mechanisms, exocyst, Rab and SNARE proteins mediate docking and fusion of secretory vesicles, ensuring the specificity of fusion between particular vesicle types and their appropriate target membranes (Saito and Ueda, 2009). These candidate regulators of *g*_c_ have potential functions in targeted delivery, to the outer face of epidermal cells, of cuticular lipids as well as proteins such as lipid transporters that are needed for cuticle assembly, thereby impacting the water barrier function of the cuticle.

The final candidate with a predicted membrane trafficking function, ISTL1, is of particular interest because it was also identified in our prior GWAS (Lin et al., 2020), and is the only candidate in the present study identified by all three approaches: GWAS, TWAS, and FCT. Yeast IST1 promotes the formation of vesicles inside multivesicular bodies (MVBs; (Hill and Babst, 2012)), a function shared by Arabidopsis ISTL1 (Buono et al., 2016). The MVB is a late endosomal compartment where plasma membrane-derived proteins are degraded after internalization via endocytosis (Lodish et al., 2016). Recent work has demonstrated that Arabidopsis ISTL1, in combination with a functionally related protein LIP5, is needed to maintain an ABCG family transporter at the plasma membrane of anther tapetal cells, and promotes the accumulation of certain wax components deposited on pollen grains by tapetal cells (Goodman et al., 2021). Thus, it can be speculated that maize ISTL1 facilitates the localization of plasma membrane transporters that export cuticle components in maize epidermal cells. Alternatively, cuticular lipids may be delivered extracellularly by fusion of MVBs with the plasma membrane.

The association between *g*_c_ and the *ISTL1* genomic region was further verified in a haplotype-based association analysis, which has increased statistical power for detecting marker-trait associations relative to GWAS, and can also identify favorable haplotypes for breeding projects. Haplotype-based association tests were performed in both the diversity panel and among nNILs. The strong correlation for haplotype effects for Block 52 in the diversity panel and nNILs confirmed the genetic associations across genetic materials and environments (2018 vs 2020). However, the power of our nNIL analysis to detect genetic effects of haplotype variation in the *ISTL1* region was reduced by two limitations. First, we analyzed only 12 nNILs, which represented all the genetic variation in this collection (Morales et al., 2020) with non-B73 haplotype in the *ISTL1* region. This is a relatively small population for this type of analysis when investigating a locus with a moderate genetic effect for a polygenic trait. In addition, Block 65, which showed the strongest association with *g*_c_ in the diversity panel, was monomorphic among nNILs, so the phenotypic effects of this haploblock could not be investigated in the nNIL analysis. While these experiments provided modest additional evidence of the impact of haplotype variation at *ISTL1* on *g*_c_, a single gene mutation analysis would be ultimately needed to prove the role of this gene.

In summary, GWAS, TWAS and FCT together revealed 22 genes and regulatory factors associated with maize adult leaf *g*_c_. These genes have predicted functions related to the synthesis and function of the cuticle. A candidate causal gene encoding ISTL1 was associated with *g*_c_ by all three tests. Haplotype-based association analysis using maize nNILs containing introgressions of the *ISTL1* genomic region from eight inbred donors into a common background (B73) corroborated marker-trait associations found in WiDiv lines, providing additional evidence of a role for *ISTL1* as a determinant of *g*_c_. The *ISTL1* region haplotypes harbored by Tzi8 were consistently associated with reduced *g*_c_, and could be introgressed into adapted maize germplasm to potentially increase drought tolerance via a reduction in non-stomatal transpiration (evaporation across the cuticle).

## Materials and Methods

### Plant materials and experimental design

A set of 323 maize inbred lines from the Wisconsin Diversity (WiDiv) panel (Hansey et al., 2011) was planted as a single replicate in a 17 × 19 incomplete block design at two different times (May and June 2018) in the same field (WiDiv18) at University of California San Diego, San Diego, CA, for evaluation of adult leaf cuticular conductance (*g*_c_). The set consisted of 89 lines that had been previously shown to have the highest or lowest *g*_c_ (extreme *g*_c_ lines) in San Diego field experiments (Lin et al., 2020) in addition to 234 lines with restricted flowering time selected for maximal genetic diversity using the CDmean method (Rincent et al., 2012). The field design in both environments was augmented by including Mo17 within each incomplete block. Experimental units were one-row plots of 3.66 m in length having approximately 12 plants with 1.02 m inter-row spacing and a 0.61 m alley. ***g*_c_ evaluation and analysis**

To inform sampling times for each plot in the WiDiv18 experiment, flowering time (days to anthesis, DTA) was scored as previously described (Lin et al., 2020). The method of Lin et al. (2020) was used to evaluate *g*_c_ on an adult leaf from the primary ear node (or one leaf immediately above/below) of five plants (no fewer than two if five were unavailable) from each plot at flowering. The calculation of adult leaf cuticular conductance (*g*_c_) from unit surface area was as follows:

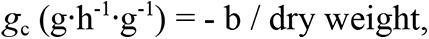

where b (g·h^-1^) is the coefficient of the linear regression of leaf wet weight (g) on time (h), and dry weight (g).

For each phenotype (*g*_c_ or DTA), a mixed linear model was fitted in ASReml-R version 3.0 (Gilmour et al., 2009) that had grand mean and check as fixed effects and environment, genotype, genotype-by-environment (G×E) interaction, incomplete block within environment, and column within environment as random effects. Model generated Studentized deleted residuals (Neter et al., 1996) were used to remove significant outliers (Bonferroni α = 0.05). The model was refitted for each outlier-screened phenotype to generate variance components to calculate heritability on a line-mean basis according to Lin et al. (2020). An iterative mixed model fitting procedure in ASReml-R version 3.0 was then used to select a best-fit model for each outlier-screened dataset (Supplemental Table S10) to generate a best linear unbiased predictor (BLUP) for *g*_c_ and DTA for each line (Supplemental Table S11) following the approach of Lin et al. (2020).

### Phenotypic and statistical analysis of extreme *g*_c_ lines

Of the extreme *g*_c_ lines evaluated in the WiDiv18 experiment, we eliminated six classified as flint, popcorn, or tropical, resulting in 51 lines mostly having membership in the NSS or SSS subpopulations (Lin et al., 2020) that were examined via super-resolution confocal microscopy. For each line, 2-5 plants from each environment were analyzed by visualizing the cuticle with lipid stain Fluorol Yellow, and images for multiple stomata were collected for each plant (*n* = 2-16 stomata per plant). Tissue samples excised from the middle portion of adult leaf blades were collected and processed with Fluorol Yellow staining to visualize cuticle features as described by Matschi et al. (2020). Collected images were processed through superresolution Airyscan, and composite pictures were processed through ImageJ. For stomatal cuticle analysis, stomata were chosen where the section presented the middle third of a stomate (where the cross sectional area of subsidiary cells was at least equal to that of the guard cells). Using ImageJ, four cuticle phenotypic features were calculated as illustrated in Supplemental Figure S1: bumpiness, cuticular flap length, roundness, and sealability. For each of the four cuticle features, means of stomatal measurements were calculated for each individual plant. Roundness was not statistically analyzed because of its very low genetic variation.

For the same 51 lines, waxes were extracted by submerging the mature leaf tissue in pure chloroform for 60 s. Extracted leaf wax samples were evaporated under a gentle stream of nitrogen, and wax samples were analyzed with gas chromatography as described in Bourgault et al. (2020).

Random forest regression models were fit to assess the prediction accuracy of *g*_c_ using 62 wax and three cuticle features. The 65 features used in modelling were calculated as an across-environment average of values that had been screened for outliers by Studentized deleted residuals (Supplemental Table S12). Forests were grown with the ‘cforest’ function from the R package ‘party’ version 1.3-7 (Strobl et al., 2007). Fivefold cross-validation was performed 50 times to evaluate the mean predictive accuracy for *g*_c_ as described in (Lin et al., 2020). Variable importance measures were calculated using the ‘varimp’ function in the ‘party’ package. Parameters used for the number of trees grown (*ntree*=1000) and predictors sampled (*mtry*=10) were those that maximized predictive accuracy.

### Construction of genomic datasets for association analyses

#### Genotypic data

The genotype data processing and imputation approach of Wu *et al*. (2021) was implemented to generate a SNP marker set in B73 RefGen_v4 coordinates for the WiDiv panel. Briefly, a target SNP set consisting of 159,878 biallelic SNPs scored on 451 WiDiv lines with all heterozygous genotypes set to missing was generated by filtering raw genotypes (exclude singletons and doubletons, call rate ≥ 20%, heterozygosity ≤ 10%, and index of panmixia ≥ 0.8) of a 955,690 genotyping-by-sequencing (GBS) SNP set obtained from Lin et al. (2020). The reference SNP genotype set of Wu *et al*. (2021), which consisted of 14,613,169 SNPs derived from maize HapMap 3.2.1 (Bukowski et al., 2018), was imputed based on the target GBS SNP set in the 451WiDiv lines via BEAGLE v5.0 (Browning et al., 2018). The resultant set of imputed genotypes was filtered to enhance SNP quality for the 310 lines with *g*_c_ BLUP values, producing a set of 9,715,072 biallelic SNPs with MAF ≥ 5% and predicted dosage *r*^2^ (DR2) ≥ 0.80 for conducting GWAS (Supplemental Data Set 1).

#### RNA sequencing data

Leaf samples for RNA sequencing (RNA-seq) were collected from plants of the WiDiv18 experiment that had not been destructively sampled for evaluating *g*_c_. In both environments, three plants at a similar stage of development per plot were selected to collect tissue samples between 9-11 am PST from the proximal, immature and actively growing 10-30% section (developing cuticle) of the total length of the longest unexpanded adult leaf (sheath length < 2 cm). The sampled leaf sections, which were immediately frozen in liquid nitrogen, were from the second, third, or fourth fully adult leaf based on the last leaf with epicuticular wax reported by Hirsch et al. (2014). For each plot, leaf tissue samples from the three plants were ground on dry ice and equally pooled to form a composite ∼100 mg tissue sample. All samples were randomized into 96-well plates for RNA extraction with the Direct-zol-96 RNA Kit (Zymo Research, Irvine, CA). Libraries constructed with the Lexogen QuantSeq 3′ mRNA-Seq Library Kit FWD (Lexogen, Greenland, NH) were sequenced on an Illumina NextSeq 500 (Illumina, San Diego, CA) at the Genomics Facility of the Cornell Institute of Biotechnology.

Raw 3′ QuantSeq reads were cleaned by trimming Illumina adaptors, the first 12 bases, and polyA tails in accordance with Lexogen recommendations. Cleaned reads were aligned to the B73 RefGen_v4 reference genome (Jiao et al., 2017) using HiSAT2 version 2.1.0 (Kim et al., 2019) with --rna-strandness F and other default parameters. Next, counts were generated in HTSeq version 0.11.2 (Anders et al., 2015) using B73 version AGPv4.42 annotation with --format=bam, --type=gene, and --stranded=yes, followed by normalization of the count data using the DESeq2 rlog function (Love et al., 2014). All genes with a normalized count of less than or equal to zero in all samples were removed.

To verify sample provenance, SNPs were called using the 3′ QuantSeq read alignments and compared to GBS SNPs (Lin et al., 2020). SNPs calls were generated from a pileup file created using SAMTools mpileup with default parameters. The called 3′ QuantSeq SNPs were filtered for quality as previously described in (Lin et al., 2020), and they were used to calculate percent identity between each pair of 3′ QuantSeq and GBS samples. With a neighbor-joining tree constructed from a percent identity-based dissimilarity matrix in R version 3.5.1 (R core team, 2018), RNAseq samples (14 in Env1 and 7 in Env2) not clustered with their corresponding GBS samples were excluded. Dataset quality was further improved by removing RNAseq samples (18 in Env1 and 28 in Env2) with < 2 million or > 20 million reads, or an overall alignment rate < 60%. The final dataset consisted of 292 (280 unique lines) samples for Env1 and 305 (288 unique lines) samples for Env2.

A mixed linear model was used to combine transcripts that had non-zero rlog-transformed counts for at least 25% of lines in each environment. To generate BLUPs for the retained 20,018 transcripts, the fitted model was the same as that used for generating phenotypic BLUPs, with the exception that sequencing lane was also included as a random effect. To account for inferred confounders, the probabilistic estimation of expression residuals (PEER) (Stegle et al., 2010) approach was applied to the 310 line × 20,018 gene matrix of BLUP expression values. The resultant PEER values after extracting 20 learned factors were screened for outliers by Studentized deleted residuals. Transcripts with missing values in over 10% of the population after outlier removal were filtered out, producing a final set of 20,013 transcripts across 310 lines for conducting TWAS (Supplemental Data Set 2).

### Association analyses

#### GWAS

Each of the 9,715,072 SNPs was tested for an association with BLUP values of *g*_c_ ^f^rom the 310 lines with a mixed linear model (Zhang et al., 2010^)^ in the R package GAPIT version 3.0 (Lipka et al., 2012) according to Lin et al. (2020). In brief, the mixed linear model controlled for maturity, population stratification, and unequal relatedness by including DTA BLUPs, principal components (PCs) based on a genotype matrix, and a kinship matrix. The PCs were calculated from the 310 line × 9,715,072 SNP genotype matrix with the ‘prcomp’ function in R version 3.5.1 (R core team, 2018). The same SNP genotype matrix was pruned at a linkage disequilibrium (LD) threshold of *r^2^* ≤ 0.2 in PLINK version 1.09_beta5 (Purcell et al., 2007), resulting in a subset of 287,023 SNPs used to construct the kinship matrix based on the centered IBS method (Endelman and Jannink, 2012) in TASSEL 5.0 (Bradbury et al., 2007). The optimal model for GWAS selected by the Bayesian information criterion (Schwarz, 1978) included only the kinship matrix. The amount of phenotypic variation explained by a SNP was approximated with the likelihood-ratio-based *R^2^* statistic (*R^2^* _LR_) of Sun et al. (2010).

#### TWAS

With BLUP values of *g*_c_ from the 310 lines as the response variable, TWAS was performed with PEER values for each of the 20,013 expressed genes by fitting a mixed linear model using the *gwas* function with the P3D function set to FALSE in the R package ‘rrBLUP’ version 4.6 (Endelman, 2011). The optimal model included the same kinship matrix from GWAS.

#### FCT

The FCT was used to combine GWAS and TWAS results according to Kremling et al. (2019). Briefly, the top 10% of the most associated SNPs (971,508) from GWAS were assigned to the nearest gene. The *P*-values of genes not tested in TWAS were set to 1. For each gene, the paired GWAS and TWAS *P*-values were used to conduct a FCT with the sumlog method in the R package ‘metap’ version 1.4 (Dewey, 2016).

### Candidate gene identification

Given that GWAS, TWAS, and FCT have different statistical power and structure, the rankings of *P*-values for each statistical method were used to identify potential candidate genes following that of Kremling et al. (2019). To identify candidate genes from GWAS results, a set of loci was declared among the top 0.01% of SNPs associated with *g*_c_ as described by Wu et al. (2021). The search interval for candidate genes was restricted to ± 200 kb of the peak SNP for each of the declared loci following Lin et al. (2020). The top 1% of genes according to their *P*-values were selected from TWAS and FCT results, resulting in comparable numbers of candidate genes across all three methods.

### Haplotype-based association analysis of *ISTL1*

Haploblocks in a ± 200 kb region encompassing the candidate gene encoding ISTL1 were constructed using the confidence interval method (Gabriel et al., 2002) in Haploview version 4.2 (Barrett et al., 2005). The identified 91 haploblocks (Supplemental Data Set 3), which were created from an LD pruned (*r*^2^ > 0.9999) subset of 726 SNPs, were each tested for an association with *g*_c_ by fitting a mixed linear model that included the same GWAS kinship matrix in ASReml-R version 3.0 (Gilmour et al., 2009). Haplotype effects were estimated for each haploblock, and pairwise comparisons were performed using the *predictPlus.asreml* function.

### Evaluation of NILs for *ISTL1*

#### Experimental field design

To assess haplotype effects of *ISTL1* on *g*_c_ variation in a common genetic background, 13 nested near isogenic lines (nNILs) that contain introgressions harboring *ISTL1* were selected. The evaluated nNILs were derived from crosses between eight diverse inbred lines (CML69, Ki11, Ki3, NC350, NC358, Oh43, Tx303 and Tzi8) and the recurrent inbred parent B73 (Morales et al., 2020). In 2020 at the University of California San Diego, the entire experiment of 13 nNILs and their eight introgression donor parents was planted as a single replicate in a 4 × 6 incomplete block design on two different dates (June 18 and 28) in the same field. Each incomplete block was augmented by B73. Experimental units for the nNIL20 experiment were the same as those in the WiDiv18 experiment.

#### Analysis of nNILs and their ISTL1 haplotypes

The collection and processing of *g*_c_ data from the nNIL20 experiment were similar to the procedures used for the WiDiv18 experiment, with the exception that best linear unbiased estimators (BLUEs; Supplemental Table S9) were calculated. Least-squares means of nNILs and donor lines were compared to B73 with the method of Dunnett (1955) in SAS version 9.4 (SAS institute 2013).

To validate the observed strong effects of *ISTL1* haploblocks on *g*_c_ in the WiDiv panel, the haploblocks most significantly associated with *g*_c_ in the WiDiv panel were investigated among 12 of the 13 nNILs that produced plants. With HapMap 3.2.1 SNP genotypes in B73 RefGen_v4 coordinates (Bukowski et al., 2018) extracted from *ISTL1* haploblocks, haplotypes for B73 and the donor parents were constructed according to those in the WiDiv panel and projected onto their corresponding nNILs by using information from GBS SNP markers of Morales et al. (2020). The haplotype-based association analysis for *g*_c_ was performed as described above, except that the GWAS kinship matrix was not needed to control for genetic background effects among tested nNILs.

### Data availability

All raw 3′ mRNA-seq data are available from the NCBI Sequence Read Archive under BioProject PRJNA773975. Supplemental Data Sets 1-3 are available at CyVerse (https://datacommons.cyverse.org/browse/iplant/home/shared/GoreLab/dataFromPubs/Lin_ LeafCuticleTWAS_2021).

## Acknowledgements

We especially thank Akriti Bhattarai, a BTI intern student, Albert Nguyen, Lesley Saldana De Haro, Alfredo Arriola, Hiep Ha, Jessica Davis, Cameron Garland, Anasilvia Herrera Fuentes, Maria Fernanda Salcedo, and Alondra Deras at UCSD for collecting phenotypic data and tissue samples. We thank Peter Balint-Kurti at USDA-ARS Plant Science Research Unit at NC State University for providing seeds of the maize nested near-isogenic lines. We also thank Elise Withers, a BTI intern student, for preliminary evaluation of an early generation computational pipeline for conducting TWAS. This research was supported by the National Science Foundation IOS1444507.

## Supplemental information

**Supplemental Figure S1.**

Guard cell cuticle structural features analyzed for relationship to cuticular conductance. Transverse sections of formaldehyde-acetic acid-alcohol-fixed, cryosectioned leaves were stained with Fluorol Yellow to fluorescently label the cuticle and imaged via superresolution confocal microscopy as described by Matschi et al. 2020. Images were collected at a position approximately halfway from base to tip of a guard cell pair (GC, guard cells; F, cuticle flap; B, bumps lining the inner surfaces of the guard cell pore). Note the semi-complementary pattern of bumps on adjacent GC surfaces. Structural features analyzed for correlation with cuticular conductance were defined as illustrated schematically. Roundness is the ratio between the length of a straight line measuring GC height (yellow line) and a curved line tracing the inner surface of the guard cell pore without regard to bumps (red). Bumpiness is the ratio between the length of a line tracing the bumps on the GC pore surface (green line) and a curved line tracing the guard cell pore surface without regard to bumps (red). Flap length is the length of a cuticular flap (blue line). Sealability attempts to measure the capacity of adjacent GC flaps to overlap when the GC pore is closed in the presence of variability in our image collection in the degree of stomatal closure. It was measured as the difference in length between a line connecting the tips of GC flaps (orange line 1) and a line connecting the inner surfaces of adjacent guard cell pores (orange line 2).

**Supplemental Figure S2.**

Upset plot showing the number of overlapping plausible candidate genes between a genome-wide association study (GWAS), a transcriptome-wide association study (TWAS), and the Fisher’s combined test (FCT).

**Supplemental Figure S3.**

Haplotype effect analysis of Block 52 in the Wisconsin diversity (WiDiv) panel and nested near-isogenic lines (nNILs). (A) Haplotype effects of Block 52 estimated in a mixed linear model analysis of adult maize leaf cuticular conductance (*g*_c_) in the WiDiv panel. The three haplotypes (1, 5, and 7) segregating among the nNILs are in bolded font. (B) The effects of the three segregating haplotypes estimated with the nNILs. The upper triangle of the matrix shows differences between haplotype effects, whereas the lower triangle shows the significance of the differences as *P*-values. (C) Pearson’s correlation between haplotype effects estimated in the WiDiv panel and nNILs.

**Supplemental Table S1.**

Relationship of *g*_c_ to cuticular wax composition and structural features. Included in the table are importance scores from random forest regression and Pearson’s correlation coefficients between best linear unbiased predictors for *g*_c_ and averaged observations for cuticular wax composition and structural features.

**Supplemental Table S2.**

Genomic information (B73 RefGen_v4) for the top 1% genes associated with maize leaf cuticular conductance (*g*_c_) in Fisher’s combined test in the Wisconsin Diversity panel.

**Supplemental Table S3.**

Genomic information (B73 RefGen_v4) and association statistics for the top 0.01% SNPs associated with maize leaf cuticular conductance (*g*_c_) in a genome-wide association study in the Wisconsin Diversity panel.

**Supplemental Table S4.**

Genomic information (B73 RefGen_v4) for the candidate genes residing within ± 200 kb of the peak SNPs identified in the genome-wide association study in the Wisconsin diversity panel.

**Supplemental Table S5.**

Genomic information (B73 RefGen_v4) for the top 1% associated genes for leaf cuticular conductance (*g*_c_) in the transcriptome-wide association study in the Wisconsin diversity panel.

**Supplemental Table S6.**

Rice and Arabidopsis homologs of 22 plausible candidate genes identified for leaf cuticular conductance (*g*_c_) via a genome-wide association study, transcriptome-wide association analysis and Fisher’s combined test in the maize Wisconsin diversity panel.

**Supplemental Table S7.**

Statistics of associations between *g*_c_ and 91 haploblocks in the vicinity of *ISTL1* in the Wisconsin diversity panel.

**Supplemental Table S8.**

Genomic introgressions for nested near-isogenic lines (nNILs) containing the *ISTL1* genomic region. Included in the table are physical positions and flanking markers for these introgressions.

**Supplemental Table S9.**

Best linear unbiased estimator (BLUE) values of adult maize leaf cuticular conductance (*g*_c_; g·h^-1^·g^-1^) and differences of least squares means (LSM) of *g*_c_ for nested near-isogenic lines (nNILs) containing *ISTL1* introgressions and their donor parents from two environments in San Diego, CA, in 2020.

**Supplemental Table S10.**

Best fitted models used to calculate best linear unbiased predictors for adult maize leaf cuticular conductance (*g*_c_) and flowering time (days to anthesis) for two environments for the Wisconsin Diversity (WiDiv) panel, as well as best linear unbiased estimators for *g*_c_ for two environments for the nested near-isogenic lines (nNILs), according to a likelihood ratio test (α = 0.05). The asterisk (*) indicates that a random effect term was retained in the mixed linear model, whereas the ‘x’ indicates that a fitted random effect term was not significant and removed from the mixed linear model.

**Supplemental Table S11.**

Best linear unbiased predictors (BLUPs) of adult maize leaf cuticular conductance (*g*_c_; g·h^-1^·g^-1^), BLUPs of flowering time (FT; days to anthesis) for 310 maize inbred lines from combined environments in San Diego, CA, in 2018.

**Supplemental Table S12.**

Cuticular waxes and stomatal cuticle features potentially associated with cuticular conductance (*g*_c_) for lines consistently at the upper or lower end of the range for the *g*_c_ phenotype in San Diego (2016 and 2017; Lin et al. 2020). Included in the table are group classification of lines following assignments of Hansey et al. (2011), best linear unbiased predictors (BLUPs) of adult maize leaf *g*_c_ (g·h^-1^·g^-1^), extreme phenotype classification (high or low *g*_c_), and averaged abundance of cuticular waxes (free alcohols, free fatty acids, hydrocarbons, aldehydes, wax esters and alicyclics) and stomatal cuticle features (sealability, flap length and bumpiness) from two environments in San Diego in 2018.

